# Stringency of synthetic promoter sequences in *Clostridium* revealed and circumvented by tuning promoter library mutation rates

**DOI:** 10.1101/216853

**Authors:** Paweł M. Mordaka, John T. Heap

## Abstract

Collections of characterized promoters of different strengths are key resources for synthetic biology, but are not well established for many important organisms, including industrially-relevant *Clostridium* spp. When generating promoters, reporter constructs are used to measure expression, but classical fluorescent reporter proteins are oxygen-dependent and hence inactive in anaerobic bacteria like *Clostridium*. We directly compared oxygen-independent reporters of different types in *Clostridium acetobutylicum* and found that glucuronidase (GusA) from *E. coli* performed best. Using GusA, a library of synthetic promoters was first generated by a typical approach entailing complete randomization of a constitutive thiolase gene promoter (P_*thl*_) except for the consensus -35 and -10 elements. In each synthetic promoter, the chance of each degenerate position matching P_*thl*_ was 25%. Surprisingly, none of the synthetic promoters from this library were functional in *C. acetobutylicum*, even though they functioned as expected in *E. coli*. Next, instead of complete randomization, we specified lower promoter mutation rates using oligonucleotide primers synthesized using custom mixtures of nucleotides. Using these primers, two promoter libraries were constructed in which the chance of each degenerate position matching P_*thl*_ was 79% or 58%, instead of 25% as before. Synthetic promoters from these ‘stringent’ libraries functioned well in *C. acetobutylicum*, covering a wide range of strengths. The promoters functioned similarly in the distantly-related species *Clostridium sporogenes*, and allowed predictable metabolic engineering of *C. acetobutylicum* for acetoin production. Besides generating the desired promoters and demonstrating their useful properties, this work indicates an unexpected ‘stringency’ of promoter sequences in *Clostridium*, not reported previously.

**GRAPHICAL ABSTRACT:** 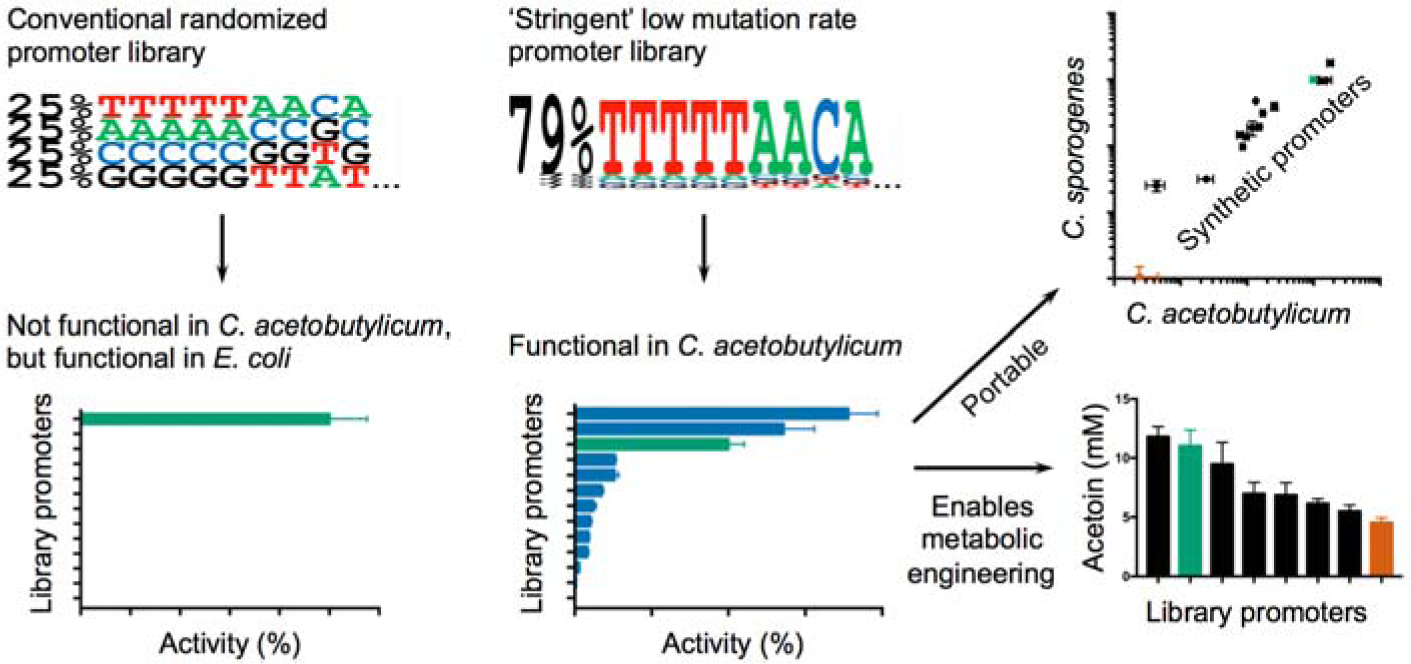

---

*Clostridium* is a large genus of anaerobic bacteria of great importance to sustainable production of chemicals and biofuels as it includes various strains that can utilize diverse feedstocks and show unique biosynthetic capabilities^1^. Moreover, *Clostridium* strains show superior tolerance to feedstocks contaminated with toxic chemicals, such as those generated by feedstock pretreatment, and to the products of fermentation^2^. *Clostridium* spp. have a long history of large-scale industrial use, and processes based on them are being developed and commercialized now, including both improved versions of historic processes and wholly new ones^1,3^. However, strain and process engineering is needed to optimize industrial-scale production of chemicals using *Clostridium* spp. For many years metabolic engineering of acetone-butanol-ethanol (ABE)-producing strains like *Clostridium acetobutylicum* had been hampered by a lack of fundamental tools allowing genetic manipulation. Recently there has been progress in the development of broadly-applicable genetic tools for *Clostridium*, including an *E. coli*-*Clostridium* shuttle plasmid system^4^, a bacterial Group II intron-based gene knockout system, ClosTron^5,6^, DNA integration by allele-coupled exchange^7^ and genome editing by allele exchange^8–11^ or CRISPR-Cas9-based systems^12,13^.

Expression control elements with known functional characteristics, including promoters and ribosome binding sites, are important to allow rational modification of native metabolic pathways and construction of synthetic and heterologous pathways. However, metabolic engineering using such elements has not been well established in *Clostridium*. Instead, protein expression has relied upon natural promoter sequences such as the *C. acetobutylicum* thiolase gene promoter^14,15^, *C. sporogenes* ferredoxin gene promoter^4^, phosphotransbutylase gene promoter^16,17^ or acetoacetate decarboxylase gene promoter^15,16,18^. However, natural promoters offer only a limited range of specific strengths and are typically subject to native regulation and variation under changing conditions^19^. In other organisms, the limitations of natural promoters have been avoided by generating synthetic promoter libraries using the approach first described by Jensen and Hammer^20,21^. In contrast to classical promoter mutants, randomization of the regions surrounding consensus -35 and -10 elements, and changing the length of the spacer between them, resulted in a library of synthetic promoters with a wide range of strengths covered in small steps, making the library suitable for fine-tuning of gene expression. This method has been applied to generate synthetic promoters for many organisms including *Lactococcus lactis*, *Corynebacterium glutamicum*, *Streptomyces coelicolor* and *Saccharomyces cerevisiae*^19^.

To generate a promoter library, reporter proteins are used to measure the strengths of individual promoter variants. However, reporter proteins are a well-known limitation for anaerobic molecular microbiology and synthetic biology. Classical fluorescent reporter proteins such as green fluorescent protein (GFP) are inactive in anaerobic bacteria, as they strictly require molecular oxygen for maturation^22^. Several colorimetric reporters have been used in *Clostridium* to determine promoter activities in the past including glucuronidase (GusA) from *E. coli*^23,24^, β-galactosidase (LacZ) from *Thermoanaerobacterium thermosulfurigenes*^14,25^, alkaline phosphatase from *Enterococcus faecalis* (PhoZ)^26^ and chloramphenicol acetyltransferase (CAT) from *Clostridium perfringens*^17,27,28^. Recently, new flavin-based oxygen-independent iLOV fluorescent proteins derived from Light-Oxygen-Voltage photoreceptor proteins^29,30^ have been successfully used in *Clostridium* such as Evoglow^31^ and PhiLOV^32^. Using iLOV proteins as reporters allows real time and *in vivo* measurements, similar to those generally achieved in aerobic biology using GFP and similar oxygen-dependent fluorescent proteins. SNAP-tags are alternative fluorescent probes based on human O-6-methylguanine-DNA methyltransferase which has been engineered to covalently bind derivatives of benzylguanine^33^. SNAP-tag fusion proteins labelled with commercially-available fluorescent dyes such as SNAP-Cell TMR-Star have been used for intracellular localization of proteins involved in sporulation of *Clostridium difficile*^34^. Multiple SNAP-tag substrates with different fluorescence properties are commercially available, which allows the labeling protocol to be optimized to avoid high background caused by green autofluorescence of *Clostridium* cells^35,36^. Reporters of gene expression remain an issue for anaerobic biology, as there are few published studies using the above, no direct comparisons to the best of our knowledge, and anecdotal reports of poor reporter performance.

In this study, we set out to develop a collection of synthetic promoters for *Clostridium*. We directly compared various oxygen-independent reporters to establish the most suitable, generated synthetic promoter libraries according to both typical and modified ‘stringent’ designs, compared promoters between distantly-related *Clostridium* strains and demonstrated their usefulness for metabolic engineering. The results provide the desired collection of characterized promoters but also suggest unexpected constraints on promoter sequences in *Clostridium*.

## RESULTS AND DISCUSSION

### Evaluation of oxygen-independent reporters favours glucuronidase

An ideal reporter should allow simple quantification of the protein product, its background activity should be absent or very low in untransformed cells, it should have a wide linear detection range and expression of the protein should not cause significant metabolic burden or toxicity. Fluorescent iLOV proteins derived from light-oxygen-voltage domains, unlike classical fluorescent proteins, do not require molecular oxygen for maturation. Therefore, they could be used for real time *in vivo* measurement of protein expression under anaerobic conditions without the need for an assay involving sample processing or labeling. Fluorescent iLOV proteins could also be used for selection of promoters of different strengths based on fluorescence intensity of colonies on agar plates. Two iLOV proteins were chosen for testing in *C. acetobutylicum*: CreiLOV from *Chlamydomonas reinhardtii*, which is brighter than other iLOV proteins^37^ and PhiLOV2.1 from *Arabidopsis thaliana* which is more photostable^38^. Genes encoding CreiLOV and PhiLOV2.1 were codon-optimized for expression in *C. acetobutylicum,* commercially synthesized and cloned under the strong constitutive thiolase gene promoter (P_*thl*_) in *E. coli*-*Clostridium* shuttle plasmid pMTL84122^27^, resulting in plasmids pPM15 and pPM16 respectively. These plasmids were introduced into *C. acetobutylicum* and intensity of green fluorescence was measured in cells harvested in the mid-exponential phase of growth. Neither iLOV reporter construct caused a detectable increase in fluorescence of *C. acetobutylicum* cells, showing that these iLOV proteins were not well expressed or not functional (**Fig. 1a**). Both iLOV constructs were shown to be functional by testing in *E. coli*, giving significantly increased fluorescence intensities almost 10-fold and 4-fold above the background level for CreiLOV and PhiLOV2.1 respectively (**Fig. 1a**). The *C. acetobutylicum* results may reflect the previously-reported high background autofluorescence of *Clostridium* cells^35,36^. Therefore, we sought an alternative reporter to measure promoter strengths. In order to allow direct comparison between SNAP-tag and glucuronidase (GusA) reporters, a gene encoding a SNAP-GusA fusion protein was designed as a bifunctional reporter and inserted into the shuttle plasmid pMTL84122, resulting in plasmid pPM4, which was introduced into *C. acetobutylicum.* Cells from the mid-exponential phase of growth were used for SNAP-tag labeling with two commercial fluorescent substrates SNAP-Cell 505-Star (compatible with standard fluorescein filter sets) and SNAP-Cell TMR-Star (for rhodamine filter sets) and the excess of unbound substrates was removed by repeated washing steps. Measurements showed that fluorescence of cells expressing SNAP-GusA increased significantly, but only 1.4- and 1.5-fold when compared to the control strain (**Fig. 1b**) giving a very narrow dynamic range of the reporter activity. The same cultures were used to quantify glucuronidase activity by a colorimetric assay. Toluene-treated cells of *C. acetobutylicum* expressing SNAP-GusA hydrolyzed the reaction substrate *p*-nitrophenyl-β-D-glucuronide forming a yellow product, which was measured using a spectrophotometer (**Fig. 1c**). A time course experiment showed a linear increase in the product concentration up to 30 min (data not shown). No glucuronidase activity was detected in the control strain expressing the empty plasmid. Therefore, GusA was found to give the highest sensitivity and the lowest background signal and was used in subsequent experiments as a reporter.

**Figure 1:**
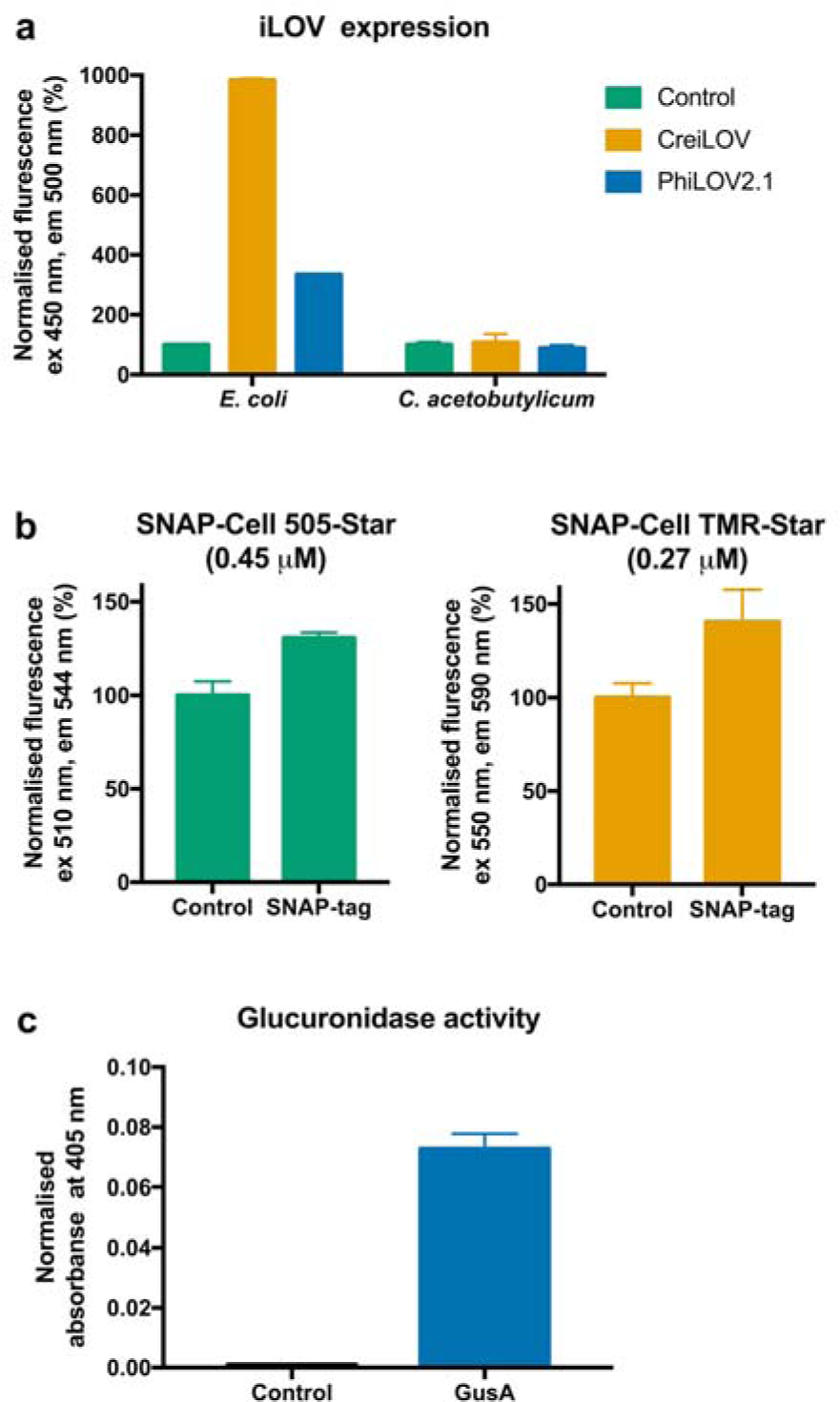
Comparison of reporter genes activity in *C. acetobutylicum*. **(a)** Expression of oxygen-independent iLOV proteins in *E. coli* NEB 5-alpha and *C. acetobutylicum*. Both strains were transformed with plasmids pPM15 and pPM16 expressing CreiLOV and PhiLOV2.1 respectively. Cells were harvested in the mid-exponential phase of growth, pelleted and resuspended in PBS. Fluorescence was measured at excitation 450 nm and emission 500 nm using using a 96-well plate monochromator and normalized to the OD of sample. Signal of the negative control (strain transformed with the empty plasmid pMTL84122) equals 100%. Error bars represent standard deviations of three independent experiments. **(b)** SNAP-tag labeling in *C. acetobutylicum*. Strains transformed with a SNAP-GusA expression plasmid were grown until mid-exponential phase. Cells were labelled with SNAP-Cell 505-Star and SNAP-Cell TMR-Star fluorescent substrates and excess of substrates was removed by washing. Fluorescence was measured at excitation 510 nm and emission 544 nm for SNAP-Cell 505-Star and at excitation 550 nm and emission 590 nm for SNAP-Cell TMR-Star. Fluorescence signal was normalized to OD600 nm. Signal of the negative control (strain transformed with the empty plasmid pMTL84122) equals 100%. Error bars represent standard deviations of three independent experiments. **(c)** Glucuronidase activity in *C. acetobutylicum*. Strains transformed with a SNAP-GusA expression plasmid and a control plasmid pMTL84122 were grown until mid-exponential phase, harvested and tested in the glucuronidase assay. Absorbance at 405 nm was normalized to OD600 nm and reaction time. Error bars represent standard deviations of three independent experiments.

### Conventional randomisation yields promoters which function in *E. coli* but not *Clostridium*

We set out to construct a synthetic promoter library for the primary vegetative sigma factor of *C. acetobutylicum* (SigA) in order to allow constitutive expression of genes at different levels in *C. acetobutylicum* and hopefully other *Clostridium* spp. The promoter of the thiolase gene of *C. acetobutylicum* was chosen as a template for the design because this promoter has been characterized with respect to the position of the consensus -35 and -10 elements and the transcription start site^39^, is active in both exponential and stationary growth phases of *C. acetobutylicum*^40^, and has been successfully used for constitutive expression of heterologous and native genes^14,16,41,42^. The promoter library was generated using the approach described by Jensen and Hammer^20^ by randomizing 39 bases surrounding the -35 (TTGATA) and -10 (TATAAT) elements in P_*thl*_ (16 bases upstream of the -35 element, the 17-base spacer between the -35 and -10 elements and 6 bp downstream of the -10 element; **Fig. 2a**). The library was generated by inverse PCR using plasmid pPM12 (which contains *gusA* under P_*thl*_ in vector pMTL83122) as template with degenerate 5’-phosphorylated primers. These primer sequences contain only the appropriate specific nucleotide at positions corresponding to the -35 and -10 elements, but contain a mixture of nucleotides specified as ‘N’ (for ‘any Nucleotide’ in the standard IUPAC notation) at degenerate positions. This means that oligonucleotide synthesis actually results in a mixed pool of oligonucleotides with the four nucleotides A, T, G and C each incorporated at approximately the same 25% frequency at each N position. The PCR products were then circularized by ligation and the ligation products used to transform *E. coli* to obtain a ‘25%’ library of clones with different synthetic promoter variants. Colonies were picked at random and the plasmids from these, containing synthetic promoters, were sequenced. The synthetic promoters showed the expected incorporation of nucleotides at each degenerate position and on average the expected 25% identity to P_*thl*_. The number of mutations varied from 27 to 34 across the 39 bp of sequence that was randomized (**Fig. S1**). Some sequenced promoter variants contained large deletions, and these were rejected. Eleven constructs with randomized promoter sequences (plasmid set pPM36-25%) were chosen for further characterisation in *E. coli* JW1609 (Keio collection^43^ *gusA* knockout mutant, chosen to minimise background glucuronidase activity) and in *C. acetobutylicum* ATCC 824. As expected, activity assays in *E. coli* showed a wide distribution of promoter strengths (**Fig. 2b**), ranging from no detectable activity (pPM36-25%-31) to a promoter 4.4-fold stronger than P_*thl*_ (pPM36-25%-39). However, none of the same set of synthetic promoter constructs showed any detectable activity in *C. acetobutylicum* (**Fig. 2c**). To the best of our knowledge, this is the first example of a library of synthetic promoters generated in the typical way (by randomizing the promoter sequence except for core -35 and -10 elements) in which all promoters are inactive. It is interesting that all the synthetic promoters generated were active in *E. coli* but not in *Clostridium*. We hypothesize that the *E. coli* primary sigma factor RpoD may be less specific and may recognise a wider range of promoter sequences than the *C. acetobutylicum* vegetative SigA subunit, which appears to be more stringent.

**Figure 2:**
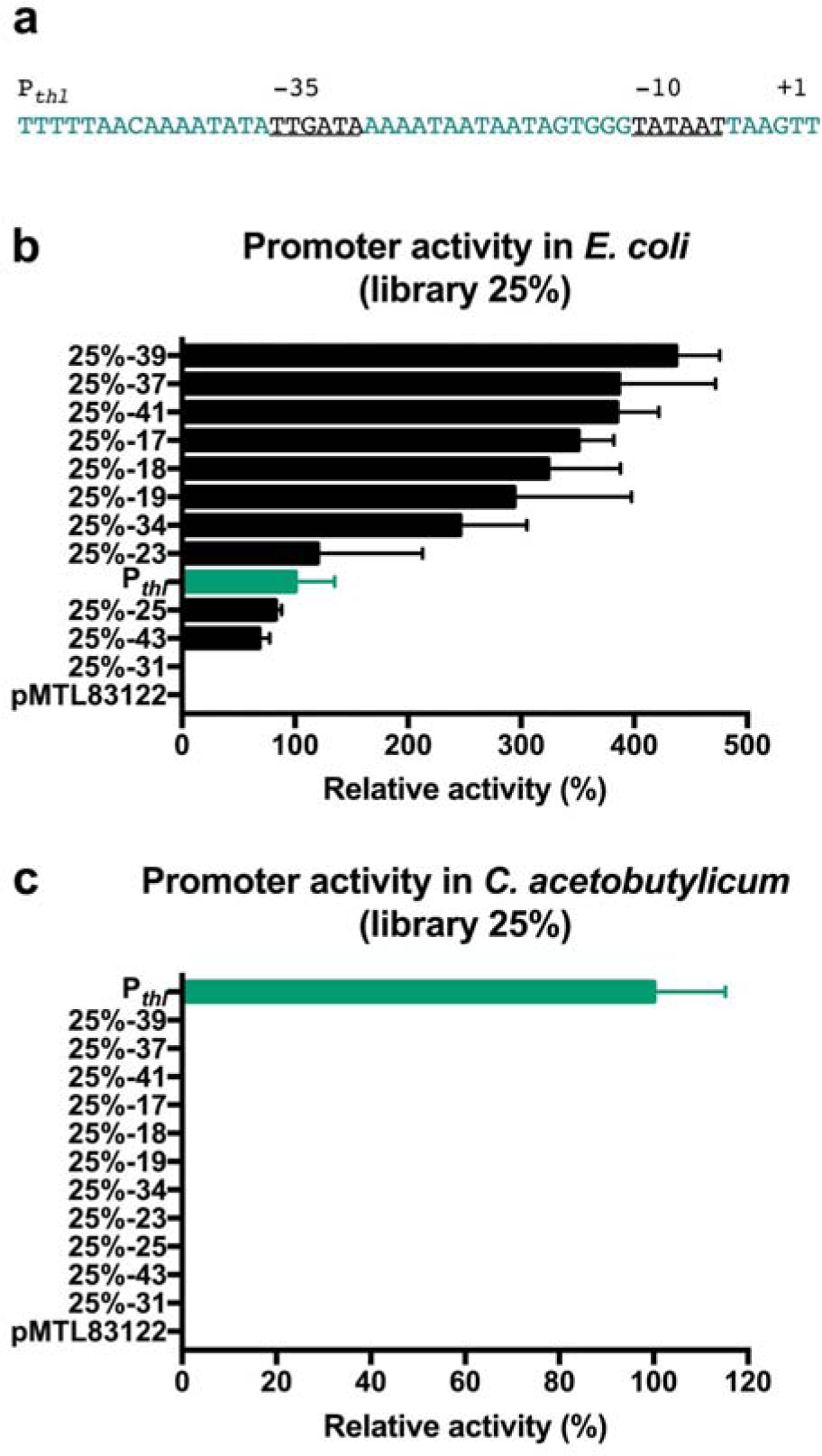
(a) Sequence of the thiolase gene promoter (P_*thl*_). -35 and -10 elements are underlined and colored black. Green bases were randomized for generation of synthetic promoter libraries. Conventional synthetic promoter library in *E. coli* JW1609 **(b)** and *C. acetobutylicum* ATCC 824 **(c)**. Glucuronidase activity of mid-exponential phase cultures harboring plasmids of the pPM36-25% library was normalized to the activity of positive control strain transformed with plasmid pPM36-Pthl (*gusA* under P_*thl*_). The same strain harboring an empty plasmid pMTL83122 was used as a negative control. Error bars represent standard deviations of three independent experiments.

### Manually tuning mutation rates yields functional promoters with diverse strengths in *Clostridium*

All tested synthetic promoters generated using the conventional 25% promoter library design described above were inactive in *C. acetobutylicum*. Presumably the sequence space of this library design does contain functional promoters besides the parent P_*thl*_, but at a low frequency. To identify a reasonable number of functional promoters, screening of a much larger number of candidates would seem to be required, ideally by visual identification of colonies expressing the reporter protein on agar plates. However, the GusA glucuronidase reporter system used in the experiment does not allow for high-throughput screening of active promoters, as blue-white colony screening using the glucuronidase substrate X-Gluc requires molecular oxygen for the development of a blue color and so cannot be used in anaerobic bacteria^26^. Both to avoid the need for laborious screening, and to investigate the apparent inflexibility of sequences of synthetic promoters in *Clostridium*, we set out to generate libraries of promoters showing higher sequence similarity to P_*thl*_, assuming that these would be enriched for functional promoters relative to the typical ‘25%’ design used previously. One approach would be to limit the number of bases randomized, but we reasoned that this might excessively limit the range and distribution of promoter strengths. Instead, the frequency of mutations in the promoter was controlled by using oligonucleotide primers synthesized using custom mixtures of nucleotides. In degenerate positions specified as N there is a 25% chance of the base incorporated at each position matching P_*thl*_, so on average 29-30 mutations relative to P_*thl*_ would be expected over the 39 degenerate positions (**Fig. 3a**) as in the 25% library. Increasing the representation of the base matching P_*thl*_ at each degenerate position would result in more conserved promoter libraries, with fewer mutations relative to P_*thl*_. For this purpose, primers were designed with custom mixtures of bases at degenerate positions corresponding to either 79% frequency or 58% frequency of the base matching P_*thl*_ at each position (and 7% or 14% frequencies for each of the three remaining nucleotides, respectively) in order to generate synthetic promoters with on average 8 or 16 mutations relative to P_*thl*_ over the 39 degenerate positions, respectively (**Fig. 3a**). These primers were synthesized and used in the same procedure as before to generate ‘79%’ and ‘58%’ promoter libraries. As before, *E. coli* colonies were picked at random, the plasmids were sequenced, and faulty promoters (with large deletions) were rejected. Eleven synthetic promoters from each library without deletions were evaluated. The average number of mutations in the 79% library (plasmids pPM36-79%) and 58% library (plasmids pPM36-58%) were 8 and 18 respectively (**Fig. S1**) which was consistent with expectations (**Fig. 3a**). Reporter constructs were introduced into *C. acetobutylicum* and promoter activities were measured. All eleven synthetic promoters from the 79% library were functional in *Clostridium* (**Fig. 3b**) showing a wide distribution of strengths (more than 260-fold change between the strongest and the weakest promoters). In the 58% library only three promoters showed detectable activities (**Fig. 3c**). In both libraries the strongest promoters were stronger than the parental promoter P_*thl*_. In *C. acetobutylicum*, unlike in *E. coli* (**Fig. S2**), the greater the number of mutations in a synthetic promoter with respect to P_*thl*_, the weaker its activity (**Fig. 4**). However, some exceptions were observed, such as synthetic promoter 58%-20, which had 17 mutations, but showed 1.55-fold higher activity than P_*thl*_. This suggests that some positions in the promoter sequence are probably more important for interaction with RNA polymerase than others and that less important bases can be mutated without inactivating the promoter. Thus, a collection of synthetic promoters (**Fig. S1**) which are functional and show a wide range of strengths in *C. acetobutylicum*, covered in small steps between promoters, was successfully generated by tuning promoter mutation rates during construction of the library.

**Figure 3:**
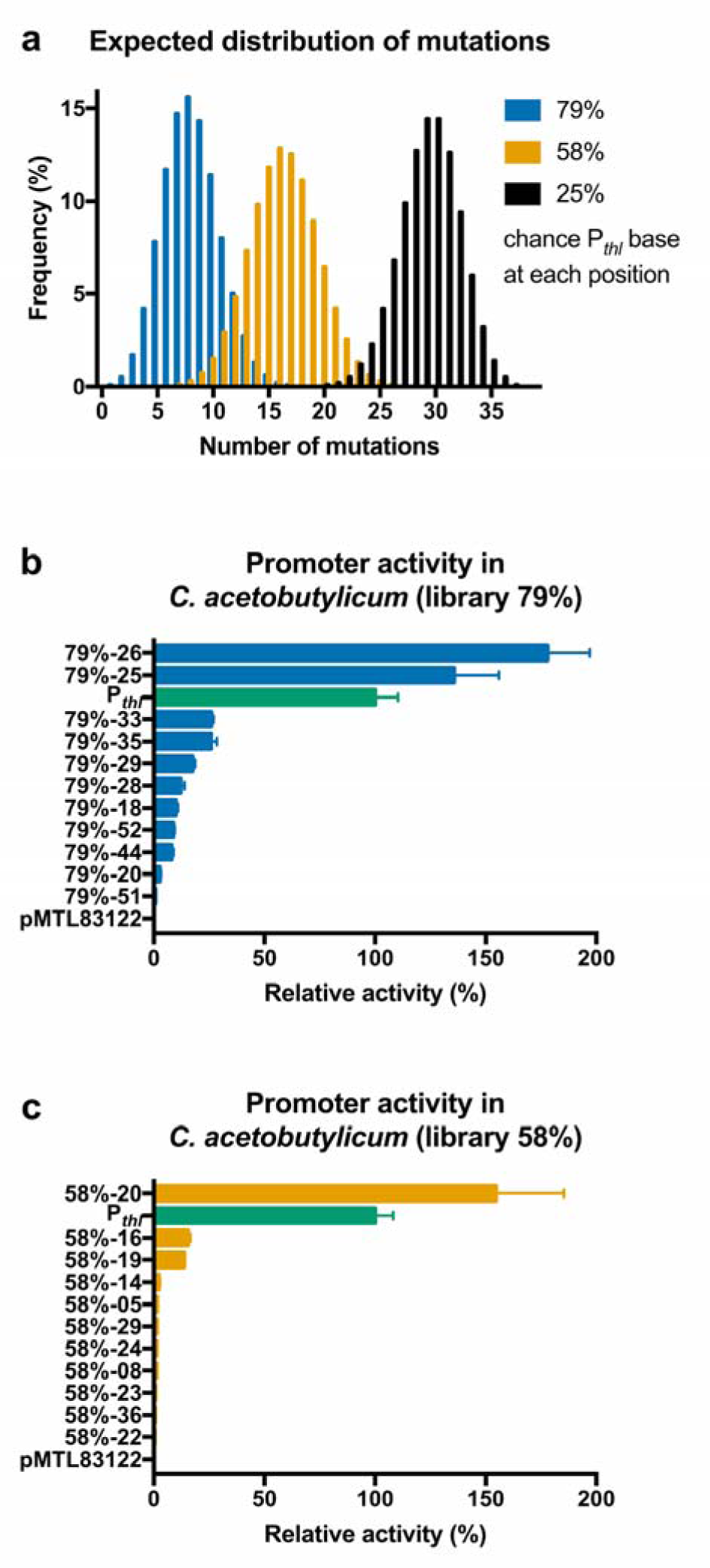
Custom mixed base synthetic promoter libraries in *C. acetobutylicum*. **(a)** Expected distribution of number of mutations in a 39 bp of degenerate sequence depending on representation of bases in the custom mixed base primer. Frequency was calculated using the binomial probability mass function. **(b)** Glucuronidase activity of mid-exponential phase cultures harboring plasmids of pPM36-79% and **(c)** pPM36-58% libraries. Activity was normalized to the activity of a positive control strain transformed with plasmid pPM36-Pthl (*gusA* under the P_*thl*_). Strain harboring an empty plasmid pMTL83122 was used as a negative control. Error bars represent standard deviations of three independent experiments.

**Figure 4:**
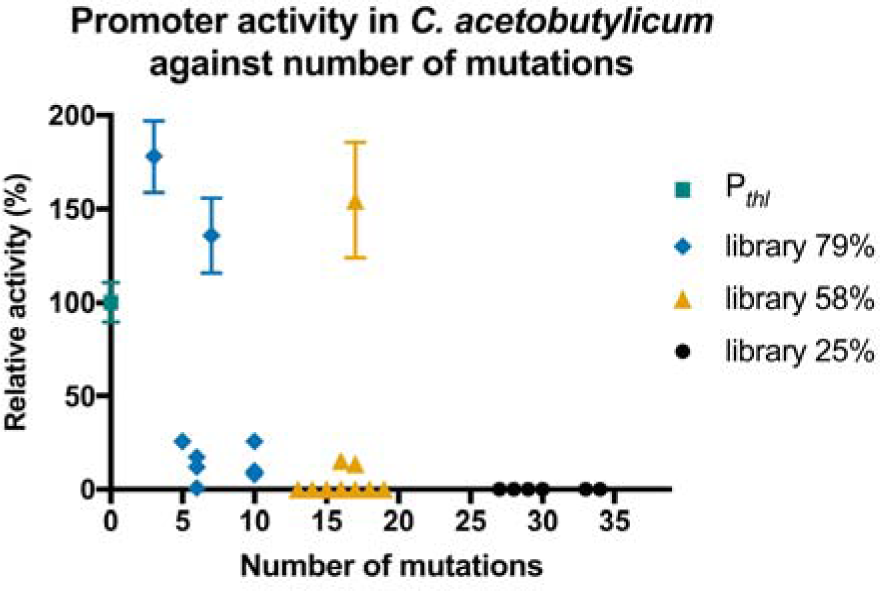
Relative activity of synthetic promoters in *C. acetobutylicum* against number of mutations. P_*thl*_ was used as a positive control. Error bars represent standard deviations of three independent experiments.

### Expression of acetolactate decarboxylase in *C. acetobutylicum* using synthetic promoters

To demonstrate the usefulness of the collection of synthetic promoters, they were applied to metabolic engineering of *C. acetobutylicum* for acetoin production. Acetoin is an important chemical building block and food flavouring, and a minor product of ABE fermentation of *C. acetobutylicum*^44^ (**Fig. 5a**). Transcriptional analysis of the acetoin pathway genes in *C. acetobutylicum* B3 has shown that the limiting step of acetoin biosynthesis is decarboxylation of acetolactate^45^. Overexpression of acetolactate decarboxylase (*alsD*) from a strong phosphotransbutyrylase gene promoter improved production of acetoin, but also significantly reduced both growth and glucose consumption rates^45,46^, so this may be a case where it would be useful to be able to fine-tune gene expression during strain development. We set out to evaluate the usefulness of our synthetic promoters to finely tune the expression of *alsD* to obtain different yields of acetoin. The acetolactate decarboxylase gene *alsD* from *B. subtilis* 168 was codon-optimized for expression in *C. acetobutylicum*, synthesized and cloned into *E. coli*-*Clostridium* shuttle plasmids under transcriptional control of selected synthetic promoters (79%-20, 79%-25, 79%-26, 79%-29, 79%-33 and 79%-52) to obtain the pPM62 series of plasmids. These expression plasmids were introduced into *C. acetobutylicum* and the recombinant strains were grown in P2 medium for 72 h. The concentration of acetoin in culture supernatants was quantified by GC-MS (**Fig. 5b**). Acetoin production titres across the different recombinant strains varied from 5.5 to 11.8 mM and ranked in the same order as strengths of promoters used for expression of *alsD* (with one exception, the P_*thl*_ and 79%-25 promoters).

**Figure 5:**
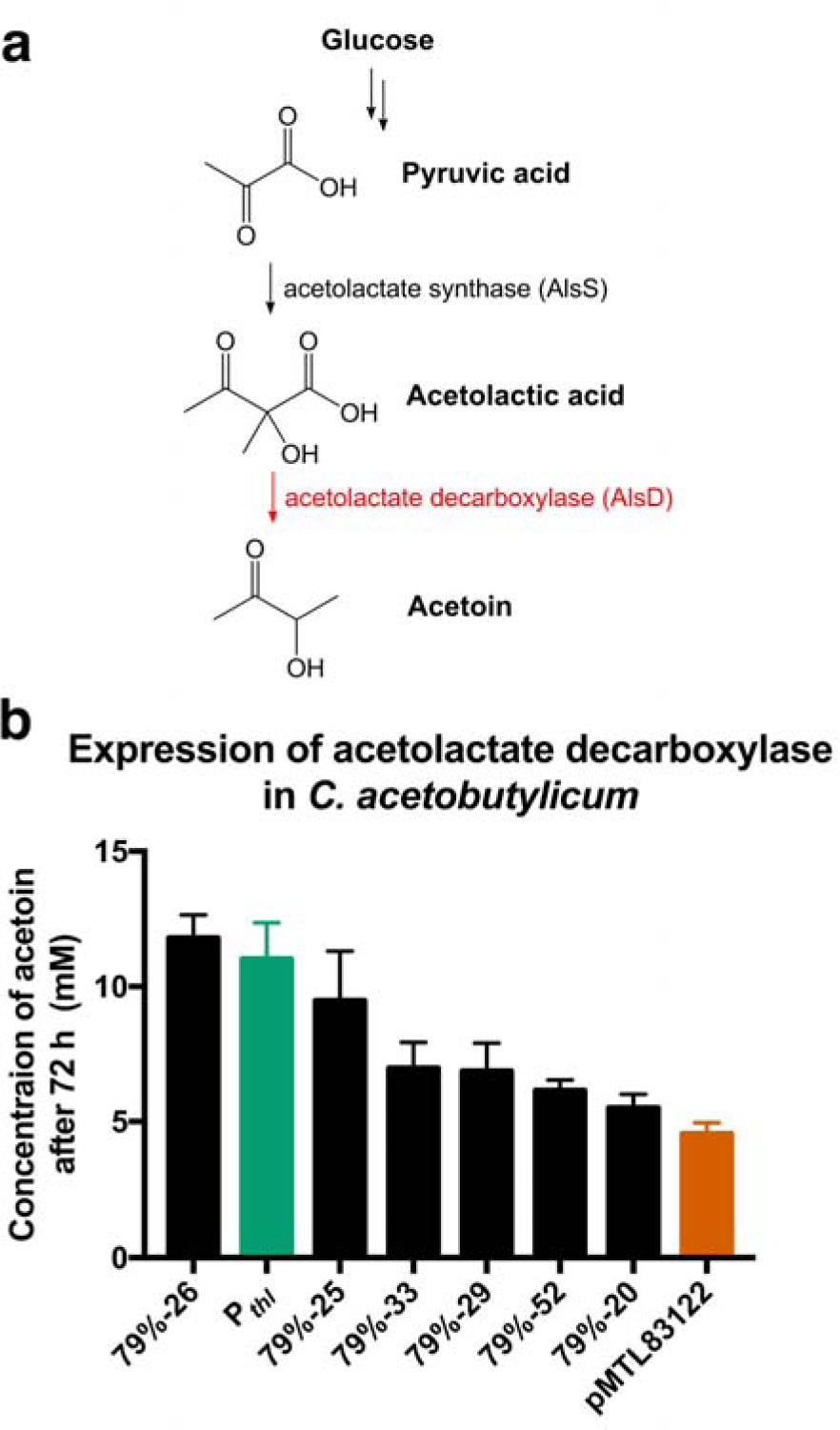
**(a)** Acetoin pathway in *C. acetobutylicum*. The limiting step (shown in red) is decarboxylation of acetolactate. **(b)** Recombinant expression of acetolactate decarboxylase from *B. subtilis* 168 in *C. acetobutylicum* ATCC 824 using synthetic promoters. *C. acetobutylicum* was transformed with pPM62 plasmid set encoding *alsD* under P_*thl*_ or selected synthetic promoters. A strain harboring an empty plasmid pMTL83122 was used as a negative control. Strains were grown in P2 medium for 72 h. Concentration of acetoin in the culture supernatant was quantified using GC-MS. Error bars represent standard deviations of three independent experiments.

### Synthetic promoters are portable between *Clostridium* spp

To test whether synthetic promoters generated in *C. acetobutylicum* are portable between different *Clostridium* species and can be used for engineering of other strains, all fourteen promoters showing activity in *C. acetobutylicum* from the pPM36-79% and pPM36-58% libraries as well as the control plasmids were introduced *via* conjugation into a proteolytic strain *Clostridium sporogenes* NCIMB 10696. This organism is of interest as a potential antitumor agent^47^ and as a safe, nontoxigenic surrogate of neurotoxin-producing *Clostridium botulinum*. Glucuronidase assays in *C. sporogenes* showed a wide range of strengths of synthetic promoters (70-fold change between the strongest and the weakest promoters) and a good linear correlation between promoter strengths in *C. acetobutylicum* and *C. sporogenes* (**Fig. 6**) with coefficient of determination R^2^ of 0.897. This data suggests that once the stringency of promoters is overcome during generation of the library, synthetic promoters are portable even between distantly related strains like *C. acetobutylicum* and *C. sporogenes*^48^.

**Figure 6:**
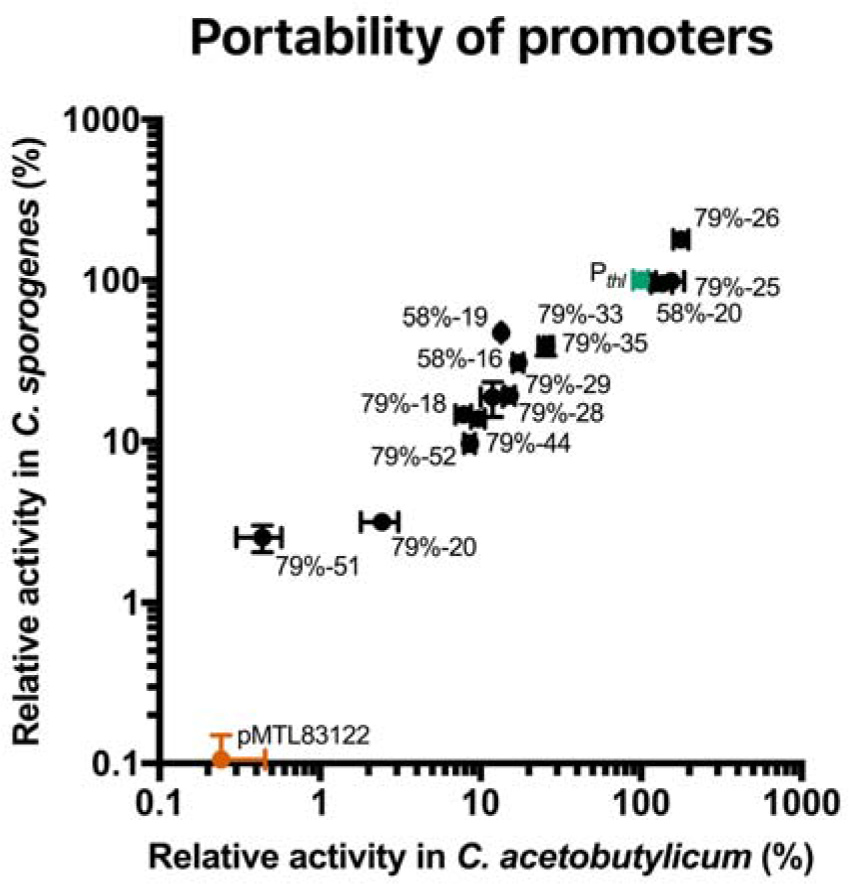
Portability of synthetic promoters between *C. acetobutylicum* ATCC 824 and *C. sporogenes* NCIMB 10696. Plasmids encoding promoters active in *C. acetobutylicum* were selected from pPM36-79% and pPM36-58% libraries and transferred to *C. sporogenes* by conjugation. Glucuronidase activity of mid-exponential phase cells was normalized to the activity of positive control strain transformed with plasmid pPM36-Pthl (*gusA* under P_*thl*_) and plotted against activity of each promoter in *C. acetobutylicum* (**Fig. 3**). Strain harboring an empty plasmid pMTL83122 was used as a negative control. Data is presented using a logarithmic scale. Error bars represent standard deviations of three independent experiments.

## CONCLUSIONS

A synthetic promoter library using the typical design, completely randomized except for the -35 and - 10 consensus sequences, resulted in promoters active in *E. coli*, but not in *C. acetobutylicum* (**Fig. 2**). This apparent stringency of functional promoter sequences in *C. acetobutylicum* was surprising, and has not been reported previously. In addition to the vegetative sigma factor SigA, *Clostridium* genomes encode multiple other sigma factors (SigH, SigF, SigE, SigG and SigK) which are central to the carefully-controlled regulatory cascade required for sporulation^49,50^. The SigA vegetative sigma factor in *C. acetobutylicum* shows only 58% identity and 76% similarity to the *E. coli* primary sigma factor RpoD^51^. We speculate that more stringent primary sigma factors in *Clostridium* recognising only specific promoter sequences might have evolved to minimise undesired crosstalk between the many different sigma factors and thereby ensure the correct cell growth and development program. Specialized sigma factors are used to control important physiological functions in many organisms^52^ so similar constraints on functional promoter sequences might exist in these. Alternatively, the apparent stringency of functional promoter sequences might be limited to organisms with particularly large numbers of alternative sigma factors, such as endospore-forming Firmicutes like *Clostridium*^49,50,53^. To the best of our knowledge the apparent stringency of promoter sequences we describe has not been reported in other endospore-forming organisms, perhaps reflecting a lack of studies of synthetic promoters that might reveal such constraints. Interestingly, another unusual constraint on design of expression constructs for *Clostridium* has recently been proposed, involving a requirement for short stem loops in 5’-UTRs for efficient expression^17^. Like the present study, this observation requires further investigation in order to establish a clear mechanism or design principles.

The apparent stringency of promoter sequence was overcome by tuning the mutation rate relative to P_*thl*_ using primers with custom mixtures of bases at degenerate positions, making the 79% and 58% libraries more conserved relative to P_*thl*_, and enriching them for promoters active in *C. acetobutylicum*. The smaller number of mutations might have been expected to narrow the range of promoter strengths, but the generated libraries showed a good distribution of promoter strengths over a 260-fold range. Other approaches could be used similarly to introduce mutations at different rates, such as error-prone PCR^54–56^ or nucleotide analogue mutagenesis^57,58^.

Exceptions to the general relationship between number of mutations and promoter activity (**Fig. 4**) suggest that some positions in the promoter sequence may be more important than others. However, if this is the case, such positions are not clear from our data set (**Fig. S1**). A much larger number of synthetic promoters with appropriate experimental design and analysis could be used in further work to investigate possible specific sequence determinants of promoter function.

During preparation of this manuscript another study describing generation of a synthetic promoter library for *C. acetobutylicum* was published by Yang and coworkers^59^. As in the first part of our study, these authors nominally used P_*thl*_ as a template and randomized most of the promoter sequence in the typical way (similar to our 25% library) meaning that the promoters generated are more accurately described as fully synthetic promoters with SigA-binding -35 and -10 elements similar to the consensus, rather than variants of P_*thl*_. Interestingly, the authors did not report a low frequency of functional promoters among the promoters tested, unlike our 25% library. However, this may reflect differences in experimental design, as Yang and co-workers effectively pre-selected their promoter library clones by directly coupling the promoter variants to expression of the antibiotic selection marker chloramphenicol acetyltransferase. Furthermore, the minimum selection applied was 100 µg mL^-1^ of chloramphenicol, which is quite high (7.5-15 µg mL^-1^ is routinely used for plasmid selection in *Clostridium* spp.) and probably excluded weak promoters from being selected for activity assays. Indeed the weakest promoter described showed only 10-fold lower activity than P_*thl*_ (resulting in only 14-fold difference between the weakest and the strongest promoter, in contrast to the 260-fold difference between the synthetic promoters described here). Furthermore, the strongest promoter in our libraries was almost 30% more active than the strongest promoter described by Yang and co-workers. Nevertheless, using antibiotic selection on agar plates was shown to be a very useful approach to increase the throughput of screening. In future the advantages of tuning mutation rates and antibiotic pre-selection could be combined to more quickly and easily generate libraries of synthetic promoters with wide expression ranges including both very strong and very weak promoters.

As a result of this work, and the recent work of Yang and coworkers mentioned above^59^, collections of characterized synthetic promoters for *Clostridium* are now available, similar to those which have proven useful for metabolic engineering in other organisms^55,56,60–62^, and should open the way to realise similar advantages in metabolic engineering of *Clostridium*.

## MATERIAL AND METHODS

### Bacterial strains, growth and plasmid transfer

A list of bacterial strains used in the study is included in the supplementary information (**Table S1)**. *E. coli* strains were grown in lysogeny broth (LB) at 37°C with rotary shaking at 250 rpm, or on LB agar plates. *E. coli* strains containing plasmids (**Table S2**) were cultured in LB broth or on LB plates supplemented with 12.5 µg mL^-1^ chloramphenicol. *Clostridium* spp. were grown in static culture at 37°C under an anaerobic atmosphere of N_2_:H_2_:CO_2_ (80:10:10, vol:vol:vol) in a Whitley A35 Anaerobic Workstation (Don Whitley, UK) in media prereduced overnight under the same conditions. Cultures were supplemented with 15 µg mL^−1^ thiamphenicol as required for plasmid selection and maintenance. *C. acetobutylicum* ATCC 824 was grown and maintained in *Clostridium* Basal Medium (CBM) or on CBM agar plates^63^. Plasmids (100-200 ng) were methylated *in vitro* using GpC methyltransferase M.CviPI (NEB) and transferred into *C. acetobutylicum* by electroporation as described previously^4^. *Clostridium sporogenes* NCIMB 10696 was grown in TYG medium or on TYG plates^64^. Plasmids were transferred to *C. sporogenes* by conjugation as described previously^4^.

### Plasmid construction

Oligonucleotide primers and synthetic DNA fragments (as ‘gBlock’ linear dsDNA from Integrated DNA Technologies) used for cloning are listed and described in the Supplementary information (**Tables S3** and **S4**). Expression shuttle plasmids (**Table S2**) for use in *E. coli*, *C. acetobutylicum* ATCC 824 and *C. sporogenes* NCIMB 10696 were based upon *Clostridium–E. coli* shuttle vectors pMTL83122 and pMTL84122^4^. To construct pPM4, the GusA coding sequence of plasmid pRPF185^24^ was PCR-amplified (Q5 Hot Start High-Fidelity DNA Polymerase, NEB) and assembled by SOE-PCR with a synthetic gBlock DNA fragment encoding a SNAP-tag codon-optimized for *C. acetobutylicum* using the Codon Optimization Tool (Integrated DNA Technologies). The resulting GusA-SNAP encoding PCR product was cloned into the pJET1.2 cloning vector (Thermo Fisher) by blunt-end ligation. The insert was excised with *Nde*I and *Xba*I and ligated to vector pMTL84122 cut with *Nde*I and *Nhe*I. pPM12 was constructed by amplification of the *gusA* sequence of plasmid pPM4, digestion with *Nde*I and *Nhe*I and ligation to vector pMTL83122 cut with the same enzymes. CreiLOV and phiLOV2.1-encoding sequences, codon-optimized for *C. acetobutylicum,* were custom synthesized as gBlock DNA fragments, digested with *Nde*I and *Nhe*I and ligated to pMTL84122 cut with the same enzymes to generate plasmids pPM15 and pPM16 respectively. pPM36-Pthl and plasmid libraries pPM36-25%, pPM36-58% and pPM36-79% were generated by inverse PCR (KOD Hot Start DNA Polymerase, EMD Millipore) with 5’-phosphorylated degenerated primers using pPM12 as a template, followed by addition of *Dpn*I to the PCR reaction to digest the template, then circularisation by ligation with T4 ligase. The pPM62 series of plasmids were generated by replacing *gusA* in plasmids selected from the pPM36-79% library with the synthetic *B. subtilis* 168 *alsD* sequence, codon-optimzed for *C. acetobutylicum,* using *Nde*I and *Nhe*I restriction enzymes.

### Fluorescent reporters

*E. coli* and *C. acetobutylicum* were transformed with plasmids pPM15 and pPM16 expressing oxygen-independent fluorescent proteins CreiLOV and phiLOV2.1 respectively. Anaerobic cultures were grown to mid-exponential phase of growth. Cells were pelleted by centrifugation for 10 min at 10000 x g and resuspended in PBS buffer to final OD600 nm of 1 (Eppendorf BioSpectrometer kinetic). Each sample (200 µL) was transferred into a black flat-bottom 96-well plate (Greiner). Fluorescence was measured at excitation 450 nm and emission 500 nm using a Spectra Max Gemini EM Microplate reader (Molecular Devices). Fluorescence signal was normalized to the negative control (strain transformed with the empty plasmid pMTL84122). For SNAP-tag labelling experiments, *C. acetobutylicum* transformed with pPM4 (expressing glucuronidase tagged with SNAP-tag at the N-terminus) was grown until the mid-exponential phase of growth. Samples (1 mL) were transferred into a deep-well 96-well plate (Corning) and sealed with a non-permeable sealing tape (Thermo Scientific). Cells were pelleted by centrifugation at 4000 x g for 10 min. After washing with PBS (800 µL) and centrifugation, pellets were resuspended in 50 µL of SNAP labeling substrate solution (0.45 µM SNAP-Cell 505-Star and 0.27 µM SNAP-Cell TMR-Star in PBS buffer). Samples were incubated in darkness for 30 min. To remove the excess of the substrate, cells were washed with PBS (800 µL) three times. Then, cells were resuspended in PBS, incubated in darkness for 30 min, pelleted and washed with PBS further three times. Finally, pellets were resupended in 200 µL of PBS and transferred into black flat-bottom 96-well plate (Greiner). Fluorescence was measured at excitation 510 nm and emission 544 nm for SNAP-Cell 505-Star and at excitation 550 nm and emission 590 nm for SNAP-Cell TMR-Star using POLARstar Omega platereader (BMG Labtech) at gain 2000. Fluorescence signal was divided by OD600 nm measured using the same platereader. Signal was normalized to the negative control (strain transformed with the empty plasmid pMTL84122).

### Glucuronidase assay

Glucuronidase activity in *E. coli* JW1609 and *Clostridium* spp. was determined as described by Dupuy and Sonenshein^23^. Briefly, strains transformed with *gusA* expression plasmids were grown to OD600 nm of 1 (Eppendorf BioSpectrometer kinetic). Samples (1.5 mL) were harvested by centrifugation and pellets were frozen at −80°C. Before testing, pellets were resuspended in 0.8 mL of buffer Z (60 mM Na_2_HPO_4_⋅7H_2_O, 40 mM NaH_2_PO_4_⋅H_2_O, 10 mM KCl, 1 mM MgSO_4_⋅7H_2_O, pH adjusted to 7.0, and 50 mM 2-mercaptoethanol added freshly). 0.2 mL of each sample was used for OD 600 nm measurement. To the remaining sample (0.6 mL), toluene (6 µL) was added, tubes were vortexed for 1 min and incubated on ice for 10 min. Tubes were transferred into a 37°C heating block and pre-incubated for 30 min with caps open. Reactions were started by addition of 120 µL of 6 mM *p*-nitrophenyl-β-D-glucuronide (Merck Millipore) solution in buffer Z. After incubation at 37°C, reactions were stopped by addition of 1 M Na_2_CO_3_ (300 µL) and reaction time was recorded. Cell debris was removed by centrifugation at 10000 x g for 10 min. Supernatants were transferred into polystyrene spectrophotometer cuvettes and absorbance at 405 nm was determined (Eppendorf BioSpectrometer kinetic). Relative glucuronidase activity was calculated by dividing the absorbance at 405 nm by sample OD600 nm and incubation time and normalization to the positive control sample (*gusA* under P_*thl*_).

### Acetoin fermentation and quantification

*C. acetobutylicum* ATCC 824 was transformed with *alsD* expression pPM62 plasmid series and pMTL83122 as a negative control. Pre-cultures in P2 medium^46^ were inoculated using fresh colonies from transformation plates and incubated overnight at 37°C. Main cultures (20 mL in 50 mL polypropylene conical tubes) in the same medium were started with 5% inoculums and incubated at 37°C for 72 h. Samples (1 mL) were harvested by centrifugation at 16000 x g for 10 min at 4°C. Supernatants were removed, filtered using 0.22 µm Nylon syringe filter (Restek) and stored at −80°C before analysis. Samples were extracted by adding an equal volume of ethyl acetate (500 µL) to the supernatant sample (500 µL), vortexing for 10 s and centrifuging for 5 min at 16000 x g. Organic phase (300 µL) was transferred into a sample vial containing a glass insert. Acetoin was quantified using a gas chromatograph (Agilent 7890B) equipped with a DB-624 Ultra Inert capillary column (30 m by 0.25 mm by 1.4 µm; Agilent) and a mass selective detector (Agilent 5977A). Ultra-pure helium was used as carrier gas at 0.8 mL min^−1^ flowrate. Samples (0.2 µL, split ratio 100:1) were injected at 240°C. The initial oven temperature was held at 35°C for 6 min, then increased at 10°C min^−1^ to 260°C, and held for 1 min. Retention times of acetoin were compared to the retention time of an authentic standard (Sigma).

## SUPPORTING INFORMATION

Supplementary Figures: Sequences of synthetic promoters, Promoter activity of pPM36-25% plasmid set in *E. coli* against number of mutations; Supplementary Tables: List of strains, List of plasmids, List of oligonucleotides, List of synthetic DNA.

## ABBREVIATIONS

ABE fermentation: acetone-butanol-ethanol fermentation
CBM: Clostridium Basal Medium
GC-MS: gas chromatography-mass spectrometry
GFP: green fluorescent protein
IUPAC: International Union of Pure and Applied Chemistry
LB: lysogeny broth
OD: optical density
PBS: phosphate-buffered saline
PCR: polymerase chain reaction
SOE-PCR: splicing by overlap extension PCR
UTR: untranslated region
X-Gluc: 5-bromo-4-chloro-3-indolyl-β-D-glucuronide

## AUTHOR INFORMATION

Author Contributions: JH and PM designed the study, PM conducted experiments, PM and JH prepared the manuscript.

Funding: This work was supported by the Biotechnology and Biological Sciences Research Council (grant reference number BB/M002454/1).

## CONFLICT OF INTEREST

The authors declare no competing financial interest.

## ACKNOWLEDGEMENT

The authors thank Prof. Neil Fairweather for supply of the plasmid pRPF185 and Dr. Soo Mei Chee for GC-MS analysis. The authors thank Dr. Linda Dekker, Lara Sellés Vidal, Dr. Ciarán Kelly and George Taylor for useful discussions.

